# Assembly methods for nanopore-based metagenomic sequencing: a comparative study

**DOI:** 10.1101/722405

**Authors:** Adriel Latorre-Pérez, Pascual Villalba-Bermell, Javier Pascual, Manuel Porcar, Cristina Vilanova

## Abstract

**Background:** Metagenomic sequencing has lead to the recovery of previously unexplored microbial genomes. In this sense, short-reads sequencing platforms often result in highly fragmented metagenomes, thus complicating downstream analyses. Third generation sequencing technologies, such as MinION, could lead to more contiguous assemblies due to their ability to generate long reads. Nevertheless, there is a lack of studies evaluating the suitability of the available assembly tools for this new type of data.

**Findings:** We benchmarked the ability of different short-reads and long-reads tools to assembly two different commercially available mock communities, and observed remarkable differences in the resulting assemblies depending on the software of choice. Short-reads metagenomic assemblers proved unsuitable for MinION data. Among the long-reads assemblers tested, Flye and Canu were the only ones performing well in all the datasets. These tools were able to retrieve complete individual genomes directly from the metagenome, and assembled a bacterial genome in only two contigs in the best scenario. Despite the intrinsic high error of long-reads technologies, Canu and Flye lead to high accurate assemblies (~99.4-99.8 % of accuracy). However, errors still had an impact on the prediction of biosynthetic gene clusters.

**Conclusions:** MinION metagenomic sequencing data proved sufficient for assembling low-complex microbial communities, leading to the recovery of highly complete and contiguous individual genomes. This work is the first systematic evaluation of the performance of different assembly tools on MinION data, and may help other researchers willing to use this technology to choose the most appropriate software depending on their goals. Future work is still needed in order to assess the performance of Oxford Nanopore MinION data on more complex microbiomes.

## INTRODUCTION

Metagenomic sequencing became a paradigm shift in the way we study and characterize microbial communities. This culture-independent technique based on shotgun sequencing has been applied in a broad range of biological fields, ranging from microbial ecology (Hiraoka *et al*., 2016) to evolution (Hug *et al*., 2016), or even clinical microbiology (Nutman and Marchaim, 2019). In recent years, metagenomics has also become a powerful tool for recovering individual genomes directly from complex microbiomes (Hug *et al*., 2016; Tully *et al*., 2018; Nayfach *et al*., 2019), leading to the identification and description of new relevant -and mainly unculturable-taxa with meaningful implications (Fettweis *et al*., 2019).

Illumina sequencing platforms have been the most widely used for metagenomics studies. Illumina reads are characterized by their short length (75-300 bp) and high accuracy (0.1-1 % of errors) (Goodwin *et al*., 2016). When performing *de novo* assemblies, Illumina sequences often result in highly fragmented genomes, even if sequencing is carried out from pure cultures (Goldstein *et al*., 2019; Wick *et al*., 2017). This is a consequence of the inability to correctly assemble genomic regions containing repetitive elements that are longer than read length (Goldstein *et al*., 2019). The fragmentation problem is magnified when handling with metagenomic sequences due to the existence of intergenomic repeats. Intergenomic repeats are genomic regions shared by more than one taxon present in the microbial community (Olson *et al*., 2017). It has to be noted that microbial communities often contain related species or sub-species in different -and unknown-abundances, resulting in extensive intergenomic overlaps that difficult the global assembly (Ayling *et al*., 2019; Sczyrba *et al.*, 2017).

Third generation sequencing platforms have recently emerged as a solution to resolve ambiguous repetitive regions and to improve genome contiguity. Despite the considerable error associated to these technologies (>10 %), their ability to produce long reads (up to 10-12 kb of mean read length) (Goodwin *et al.*, 2016; Nicholls *et al*., 2019) has allowed them to generate genomes with a high degree of completeness (Jayakumar and Sakakibara, 2017; Loman *et al*., 2015). Currently, the most widely used third generation technologies are Pacific Biosciences (PacBio) and Oxford Nanopore Techonologies (ONT), both based on single molecule sequencing, and therefore, PCR-free. PacBio was the first long-read technology to be established in the market (Koren *et al*., 2013). However, PacBio instruments require particular operation conditions and huge capital investments (Gonzalez-Escalona *et al*., 2019). On the other side, ONT platforms are becoming more and more popular between researchers, mainly thanks to MinION sequencers. MinION is a cost-effective (~1000$), portable sequencing platform, which enables real-time analysis pipelines (Lu *et al*., 2016). This platform has been broadly applied over the last few years, especially for testing their suitability for in-field or clinical applications (Pomerantz *et al*., 2018; Orsini *et al*., 2018), but also for sequencing complete prokaryotic and eukaryotic genomes (Loman *et al*., 2015; Wick *et al*., 2017; Deschamps *et al*., 2018; Jain *et al*., 2018) and for characterizing microbial communities (Hardegen *et al*., 2018; Benítez-Páez and Sanz, 2017).

Benchmarking is a straightforward way to evaluate genomic methodologies (i.e. DNA extraction, library preparations, etc.) and bioinformatic tools. In the metagenomic context, benchmarking studies are frequently based on mock communities. A mock community is an artificial microbial community in which the abundance of all the present microorganisms is known (Bokulich *et al*., 2016). Mock communities could be generated *in silico* (Fritz *et al*., 2019) or experimentally, as a mixture of defined DNA proportions. For *de novo* assemblies, a great effort has been made in order to benchmark all the available tools and methodologies suitable for studying microbial ecosystems via Illumina shotgun sequencing (Sczyrba *et al*., 2017; Vollmers *et al*., 2017; Nurk *et al*., 2017). Nevertheless, although there is a constant development of new softwares applicable to ONT platforms, we found that the few evaluation studies made for nanopore-based shotgun sequencing data have focused on reconstructing single bacterial genomes from isolates, but not metagenomes (Goldstein *et al*., 2019; Tyler *et al*., 2018; Sović *et al*., 2016).

In the present study, we used the data generated by Nicholls *et al*. (2019) to comprehensively assess the current state-of-art of *de novo* assembly tools suitable for MinION sequencing. For that purpose, we subsampled the sequences generated by GridION and PromethION platforms to get an output comparable to the current yield of MinION sequencers. In total, we generated 8 datasets consisting of 3 and 6 Gbps of data coming from the metagenomic sequencing of two microbial communites (ZymoBIOMICS Microbial Community Standards CS and CSII) with both GridION and PromethION. Our results show very notable differences in assembly performance among the tested tools, including those designed to work with long-reads. Nevertheless, Flye and Canu were able to retrieve highly complete and contiguous draft genomes directly from the metagenome, and work consistently in all the datasets. Despite the high error associated to long-reads technologies, these assemblers were able to return draft genomes with up to 99.85 % of accuracy. Overall, this work demonstrates the suitability of using MinION sequencing alone for assembling low-complex microbial communities, and paves the way towards the standardization of bioinformatic pipelines for long-reads sequencing data.

## METHODS

### Dataset description

Benchmarking datasets were extracted from Nicholls *et al*. (2019), and consisted of the high coverage sequencing of two individual mock communities (ZymoBIOMICS Microbial Community Standards CS Even ZRC190633 and CSII Log ZRC190842) with both GridION and PromethION platforms. The mock communities contained the same species (eight bacteria; two yeasts), but differed in the expected proportion for each microorganism. CS mock community has an equal distribution of the microorganisms (12% for each bacteria, and 2% for the yeasts), while the microbes present on CSII are distributed on a logarithmic scale, with relative abundances ranging from 89.1% to 0.000089% (Table 1). Following the nomenclature from Nicholls *et al*. (2019), we will now onwards use the terms “Even” when referring to CS mock community, and “Log” when referring to CSII.

**Table 1.**
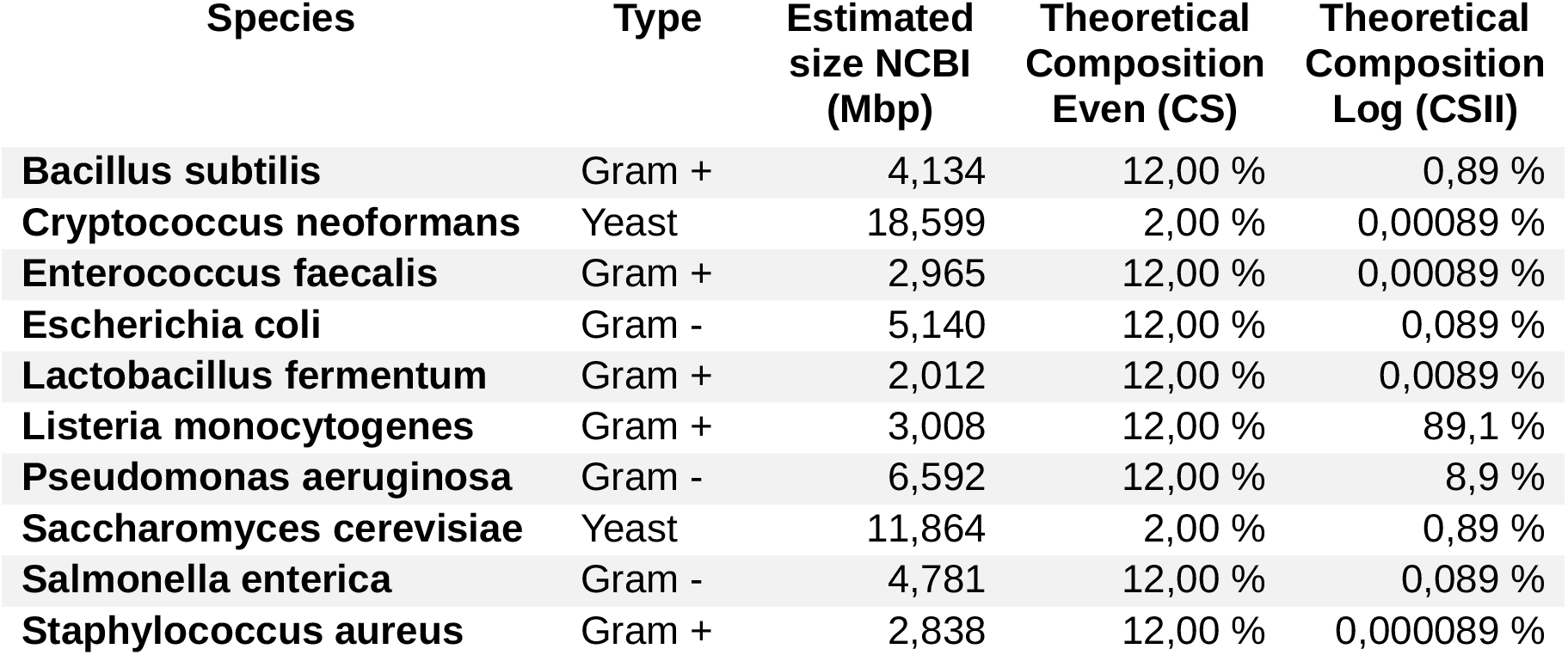
Description of the microorganisms comprising the ZymoBIOMICS mock communities and their theoretical composition.

The objective of the present study was to evaluate *de novo* assemblers suitable for MinION sequencing, which is the most widespread and accessible ONT sequencer. With the recent adoption of Guppy (Oxford Nanopore Technologies) as the lead basecaller for all the ONT sequencers, the main difference between GridION, PromethION and MinION is the final output of each platform. Nicholls *et al*. (2019) yielded ~15 Gbp of data for GridION (48h of sequencing) and ~152 Gbp for PromethION (64h of sequencing). Taking into account that GridION consists of five MinION flowcells, a single MinION standard run (48 h of sequencing) could yield, on average, an output of 3 Gbp, which is a conservative estimation in comparison to other recent shotgun sequencing experiments based on MinION (Goldstein *et al*., 2019; Dhar *et al*., 2019; Parajuli *et al*., 2019). However, ONT hardware and software are in constant development, leading to huge improvements in short periods of time. For that reason, GridION and PromethION datasets were subsampled to two different sequencing depths (3 Gbps and 6 Gbps) in order to recreate MinION runs with different outputs. Finally, all the selected reads were trimmed with porechop (https://github.com/rrwick/Porechop; v. 0.2.4) in order to remove adapters from reads ends and split sequences with internal adapters.

### *De novo* assemblers selection

As first proposed by Lindgreen *et al*. (2016), tools selected for the present benchmarking had to meet the following criteria:

- The tool should be freely available
- The tool should have a proper manual, both for installation and usage.
- The tool should have been extensively used or show potential to become widely used

At the time of the software selection, there was not a huge variety of tools specially designed for ONT data. Because of this, some of the most widespread used short-reads metagenomic assemblers were also included into the benchmark. Although these assemblers are optimized for metagenomic datasets, it has to be noted that they have not been designed to handle long and error-prone reads. A total of six short-reads and six long-reads tools were taken into consideration. Nevertheless, it was not possible to install or run all the softwares for different reasons (Table 2). It has to be noted that tools were run with default parameters when no metagenomic configuration was explicitly recommended in the user guide.

**Table 2.**
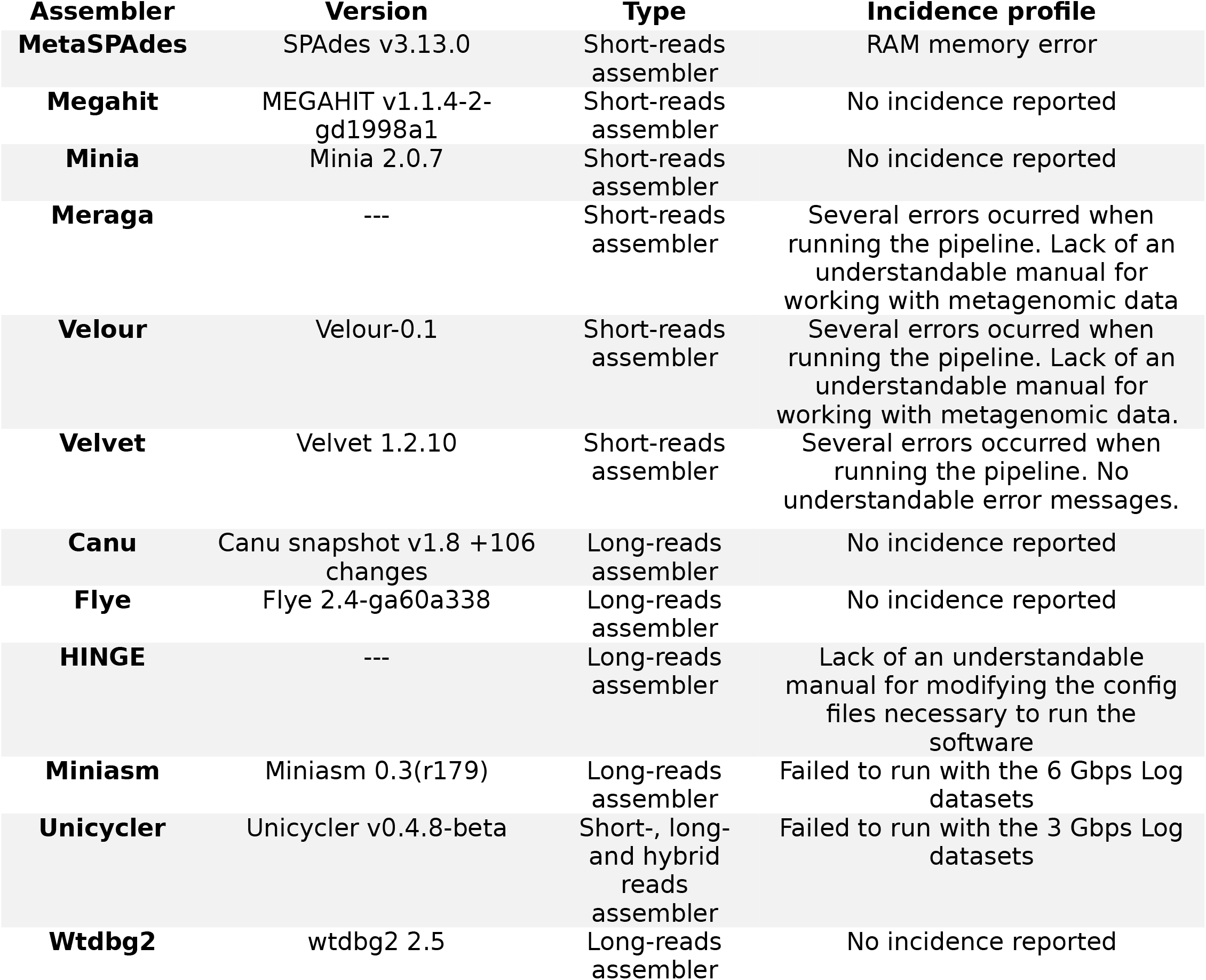
List of assemblers selected for the present benchmarking study.

### Reference genomes

All the species included in the mock community had an available reference genome sequenced with a combination of Illumina and nanopore reads (available at https://s3.amazonaws.com/zymo-files/BioPool/ZymoBIOMICS.STD.refseq.v2.zip). These assemblies -provided by ZymoBIOMICS company-consisted of eight complete genomes for the bacterial strains, and two draft genomes for the yeasts.

Nicholls *et al*. (2019) sequenced and assembled each genome again from pure cultures using Illumina reads only. However, we decided to use ZymoBIOMICS genomes as a reference for carrying out the comparative analyses, due to their higher level of completeness. Although these references cannot be considered as a “gold standard”, Goldstein *et al*. (2019) demonstrated that nanopore sequences polished with Illumina reads had a similar error profile to MiSeq-only assemblies and higher contiguity. Reference genomes were gathered in a single multi-FASTA file to create a single-reference metagenome.

### Evaluation of the assembly tools

All the assemblers were run in the same desktop computer (CPU: AMD RYZEN 7 1700X 3.4GHZ; Cores: 8; Threads: 16; RAM: Corsair Vengeance 64 GB; SSD: Samsung 860 EVO Basic SSD 500GB) working under Ubuntu 18.04 operative system. Time required to perform the assembly by each tool was measured with the built-in bash version of time command.

*De novo* assemblies completeness and contiguity were first evaluated via QUAST (Gurevich *et al*., 2013; v. 5.0.2). MetaQUAST (Mikheenko *et al*., 2015; v. 5.0.2) was used for obtaining further assembly statistics based on the alignment of the generated contigs against the reference genomes. Only contigs longer than 500 bp and with x10 coverage or more were selected for calculating the general statistics. MetaQUAST failed to run with some draft metagenomes. For that reason, minimap2 (Li *et al*., 2018; v.) was employed to align the assemblies to the reference metagenome. Then, ‘pileup.sh’ script from BBTools (sourceforge.net/projects/bbmap/, v. 2.15-r915) suite was utilized to calculate the percentage of metagenome covered by the draft assemblies.

The resulting assemblies were further evaluated in order to determine their error profile. Due to the lack of a standard methodology, SNPs and indels were ascertained using two different strategies. The first one consisted of the alignment of the contigs against the reference metagenome via minimap2. BAM files were then revised using bcftools (https://samtools.github.io/bcftools/; v. 1.9) and the in-house script ‘indels_and_snps.py’ (Supplementary File 1) was applied to quantify the variants. The second strategy was based on MuMmer4 (https://sourceforge.net/projects/mummer/files/; v. 3.23). This tool was employed to align the draft assemblies to the reference metagenome. Then, the script ‘count_SNPS_indels.pl’ from Goldstein *et al*. (2019) was utilized to calculate the final number of SNPs and INDELs. In both strategies, the number of variants were normalized to the total assembly size of each metagenome.

Biosynthetic gene clusters (BGCs) are usually formed by repetitive genetic structures hard to assemble with short-reads technologies, and long-read technologies could thus be suitable to overcome this issue. However, BGCs are also very sensitive to frameshift errors, which have been reported to frequently occur in nanopore data (Goldstein *et al*., 2019). For that reason, AntiSMASH web service (v. 5.0; Blin *et al*., 2019) was used to compare the performance on BGC prediction among the different assembly tools.

## FINDINGS

### Subsampling

In order to study the applicability of ONT to characterize low complex microbial communities, we used the data recently released by Nicholls *et al*. (2019), which consisted of the ultra-deep nanopore sequencing of two different mock communities by GridION and PromethION platforms. The mock communities were constituted by the same ten microorganisms, but in different proportions (Table 1). As we wanted to study the suitability of MinION to reconstruct individual microbial genomes from metagenomes, we subsampled the GridION and PromethION datasets to have a final output of approximately 3 Gbps and 6 Gbps, which is the current output of MinION. In general, mean read length remained the same in the subsampled datasets in comparison to the original sequencing data (Nicholls *et al*., 2019). However, read quality proved higher in the subsampled dataset, suggesting a bias towards lower qualities when the data volume increases (Table 3).

**Table 3.** Description of the original and the subsampled datasets.

|  | ORIGINAL DATASET |  |  |  |  | NEW DATASET |  |  |  |  |
| --- | --- | --- | --- | --- | --- | --- | --- | --- | --- | --- |
|  | Gb | Gbps | Number of reads | Mean read length | Mean read quality | Gb | Gbps | Number of reads | Mean read length | Mean read quality |
| Even GridION | 14 | 14.007 | 3,491,078.0 | 4,012.3 | 8.4 | 3 | 3.042 | 747,682.0 | 4,069.5 | 8.9 |
| Log GridION | 16 | 16.032 | 3,667,007.0 | 4,372.0 | 8.0 | 3 | 3.053 | 685,926.0 | 4,451.0 | 8.7 |
| Even PromethION | 146 | 146.291 | 36,527,376.0 | 4,005.0 | 7.3 | 3 | 2.979 | 748,367.0 | 3,981.0 | 8.2 |
| Log PromethION | 148 | 148.028 | 35,118,078.0 | 4,215.2 | 7.6 | 3 | 2.990 | 711,524.0 | 4,203.3 | 8.3 |
| Even GridION | 14 | 14.007 | 3,491,078.0 | 4,012.3 | 8.4 | 6 | 6.092 | 1,495,377.0 | 4,073.9 | 8.8 |
| Log GridION | 16 | 16.032 | 3,667,007.0 | 4,372.0 | 8.0 | 6 | 6.094 | 1,371,820.0 | 4,442.4 | 8.5 |
| Even PromethION | 146 | 146.291 | 36,527,376.0 | 4,005.0 | 7.3 | 6 | 5.970 | 1,496,919.0 | 3,988.8 | 8.2 |
| Log PromethION | 148 | 148.028 | 35,118,078.0 | 4,215.2 | 7.6 | 6 | 5.956 | 1,422,918.0 | 4,185.8 | 8.2 |

### Metagenome assembly

From the selected pool of available tools (Table 2), we were able to correctly install and run five out of the six long-reads assemblers, and two out of the six short-reads assemblers. In total, 58 assemblies were generated, 28 for the Even mock community and 24 for the Log community. The total size of each draft assembly and the fraction of metagenome recovered from the reference genomes were evaluated for the Even datasets in order to obtain a first view of the general tool performance.

Overall, long-reads assemblers resulted in a total assembly size closer to the theoretical size, and also recovered a largest metagenome fraction, with some exceptions (Fig. 1). Nevertheless, huge differences were detected for both metrics among the assemblers. In general, all the assemblers were far from recovering the totality of the metagenome, either in the 3 Gbps or 6 Gbps datasets (Fig. 1A). It has to be noted that metaQUAST and minimap2 results were consistent for the long-reads assemblers, but not for the short-reads assemblers, where minimap2 metric was significantly higher (Fig. 1B). The Flye assembler yielded the best assembly in terms of total metagenome size and metagenome recovery -except for the minimap2 metric-, whereas Canu proved the second best assembler for both dataset sizes. Interestingly, Unicycler and Miniasm performed relatively well for the 3 Gbps dataset, but when using 6 Gb, the final assembly did not improve significantly in the case of Miniasm, and the general performance was highly reduced for Unicycler. Wtdbg2 resulted in a poor assembly in comparison to the other long-reads tools for both the 3 Gbps and 6 Gbps datasets.

**Figure 1.**
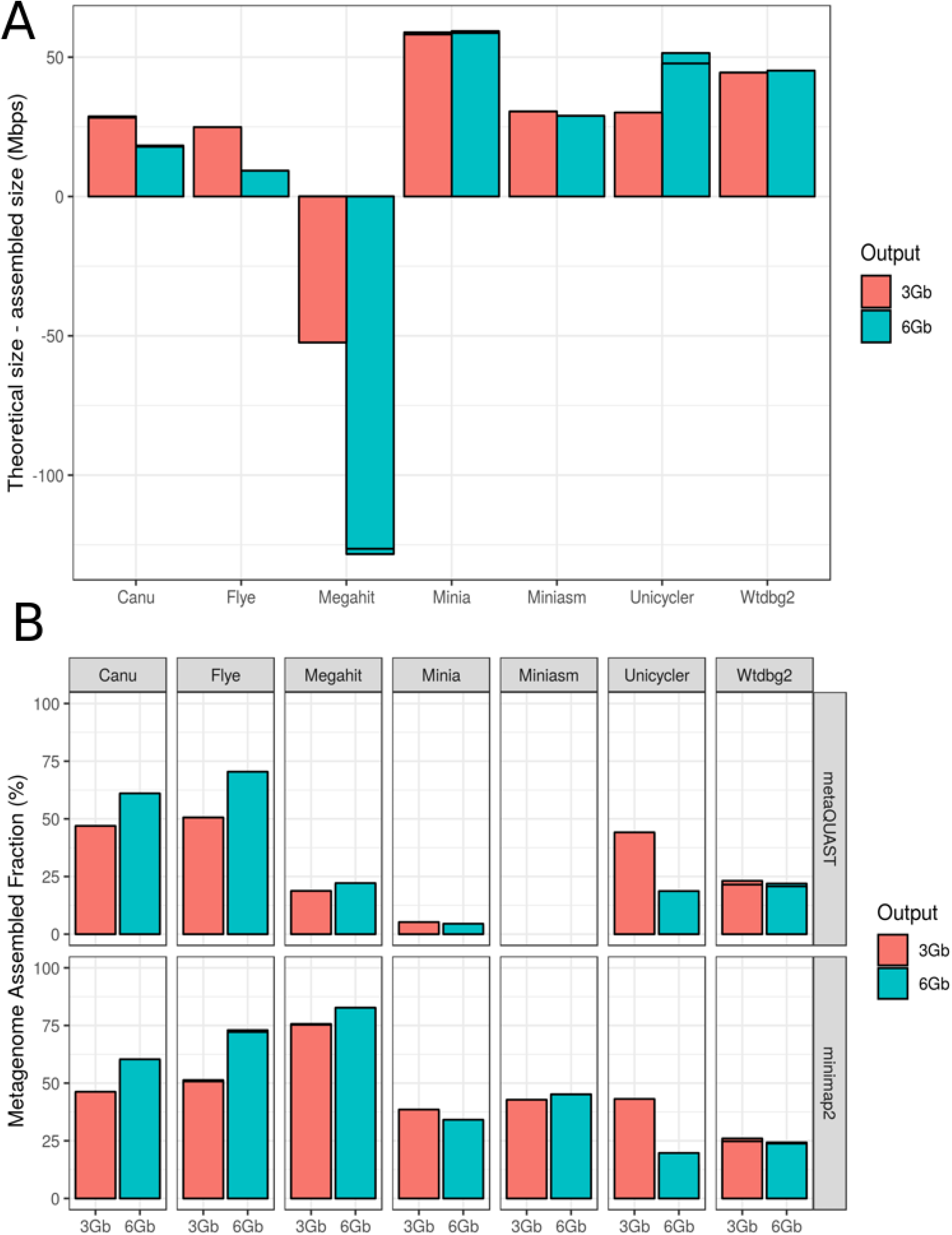
Evaluation of metagenome assembly size corresponding to each tested tool for the Even datasets. (A) Total assembled size of draft assemblies with respect to the total size of the reference metagenome; (B) Fraction of the reference metagenome covered by the draft assembly, calculated by two different methods: metaQUAST (up) and minimap2 + BBTools (down).

MetaQUAST was further employed for evaluating the degree of completeness of the individual species draft genomes (Fig. 2). As expected, yeasts were generally less recovered than bacteria, due to their lower abundance (2 %) and higher genome size. Minia and Megahit were not able to recover any single highly complete genome (>95 % of genome coverage) in any dataset, while wtdbg2 only worked well on recovering *Pseudomonas aeruginosa*’s genome. For the 3 Gbps dataset, Flye and Unicycler recovered the eight bacterial genomes with a high completeness level (> 99%). Canu resulted in lower recovery percentages, but still retrieved all the prokaryotic genomes with a mean covered fraction greater than 87%. Unicycler was able to return three totally complete genomes, but did not work properly on recovering eukaryotic genomes. This was expected, since this assembler was designed for working on bacterial genomes only. For the 6 Gbps dataset, Unicycler performance decreased substantially, while Canu and Flye retrieved better or similar results. In general, Flye performed the best on both dataset sizes, especially if taking into account the proportion of yeast genomes recovered for each tool.

**Figure 2.**
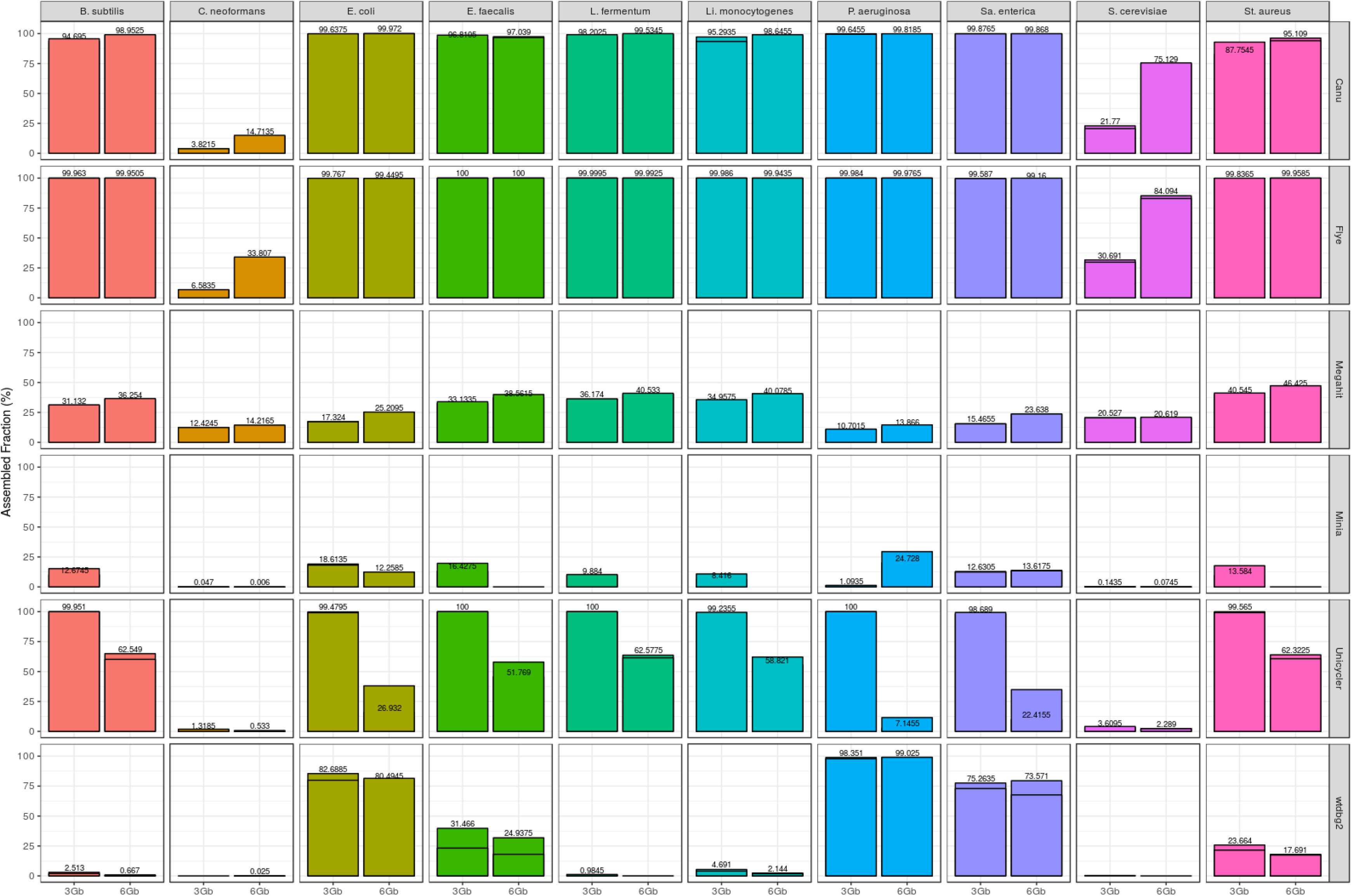
Fraction of genome covered by draft assemblies obtained with each tool, and for each individual microorganism (Even datasets). Minimap2 + miniasm assemblies are not shown, since it was not possible to evaluate them with metaQUAST.

These results were confirmed when analyzing the Log mock community (Fig. S1). Canu, Flye and wtdbg2 were able to recover *Listeria monocytogenes* (89.1% of the total genomic DNA) and *Pseudomonas aeruginosa* (8.9%) genomes with a level of completeness higher than 99%. Nevertheless, only Canu and Flye recovered a significant fraction of *Bacillus subtilis* (0.89%). Again, Flye outperformed the rest of the tools in terms of total metagenome recovery. Unicycler failed to run with the two 3 Gbps datasets, and performed poorly with the 6 Gbps ones. These results were expected, since Unicycler was designed and optimized for working with isolated bacterial genomes. Finally, short-reads assemblers resulted in highly fragmented draft metagenomes and were not able to recover any single complete genome (Fig. S1).

Regarding the time consumed by each tool, wtbdg2 was the fastest assembler (Fig. 3A). This tool was able to assemble the 6 Gbps datasets in only 155 seconds, approximately. Miniasm was the second most rapid software, followed by Flye, which was 2.1-2.5 times faster than Unicycler, and 3-5 times faster than Canu, the slowest tool. These trends were also found in the Log mock community (Fig. S2), were Canu spent up to 22 hours to reconstruct a draft metagenome assembly from the 6 Gbps datasets.

**Figure 3.**
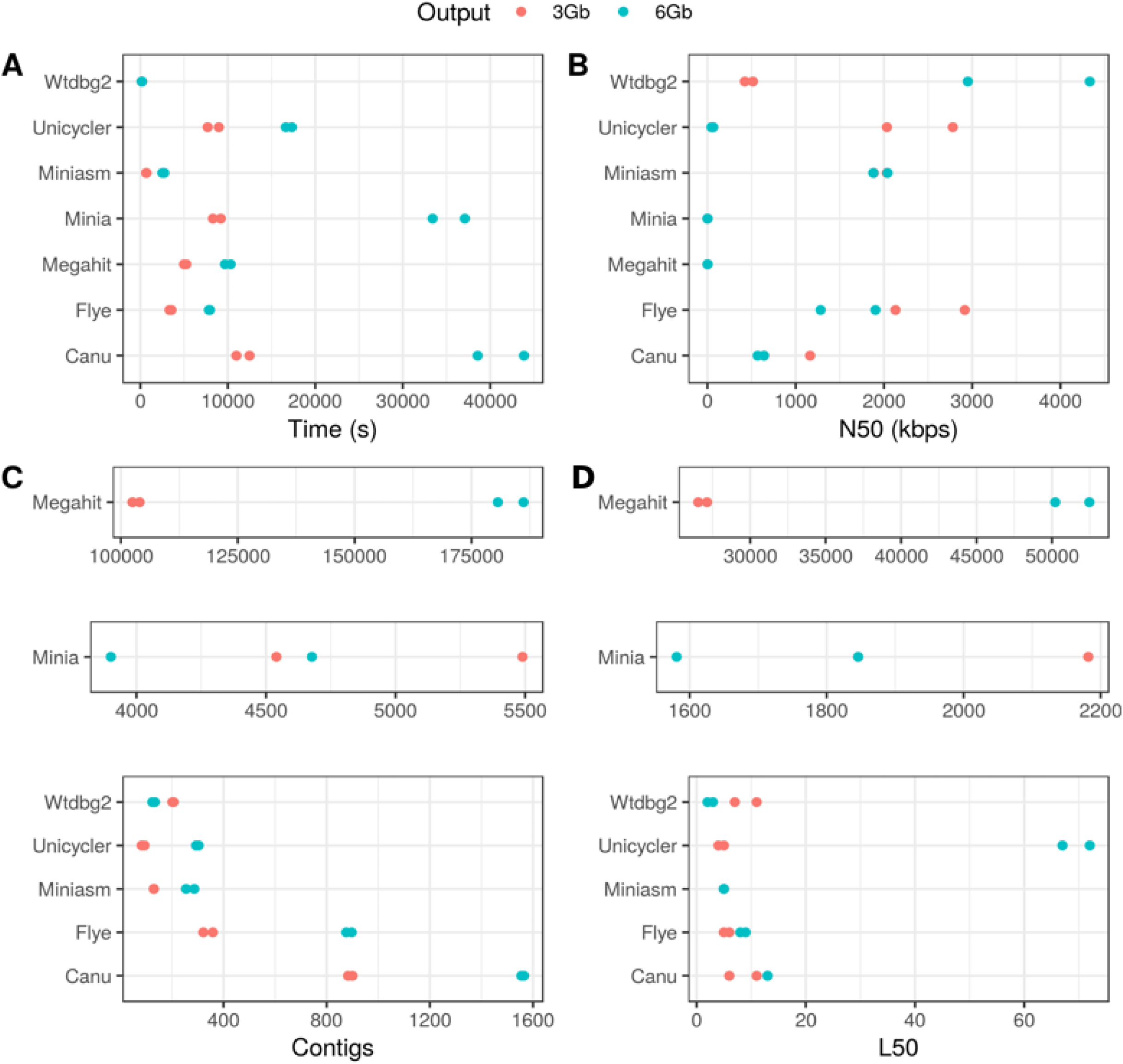
General assembly performance of each tool for the Even datasets. (A) Run time; (B) N50; (C) Number of contigs; (D) L50.

Metagenome general statistics (N50, L50, and number of contigs) were evaluated using QUAST (Fig. 3). It has to be stressed that these statistics have to be taken with care in this case, due to the huge variation in general performance among the different assemblers. For instance, wtdbg2 resulted in the higher N50 and the lower L50 values for the 6 Gbps dataset, but this tool was able to cover less than the 25 % of the metagenome. In fact, the total assembly size for wtdgb2 was approximately 18 Mbps, in comparison to the 53 Mbps assembled by Flye. Altogether, it can be concluded that N50 and L50 results for wtdgb2 were indeed an artifact.

Short-reads assemblers performed poorly, resulting in thousands (Minia), or even hundreds of thousands contigs (Megahit). Interestingly, long-reads assemblers resulted in more fragmented draft genomes when using the 6 Gbps datasets, with the only exception of wtdbg2. Flye, Canu and Unycicler also reduced their N50 and increased their L50 score when using 6 Gbps. This variation was specially marked in the case of Unicycler, confirming a worse performance of this tool when using larger datasets. Goldstein *et al*. (2019) demonstrated that Canu assemblies improved with higher coverage for bacterial isolates assemblies. This fact suggests that the loss of contiguity detected in Flye and Canu may be a direct consequence of a higher recovery rate of yeast genomes, which might be more fragmented. Indeed, assembly statistics of these two assemblers remained almost the same for the bacterial species when using 3 or 6 Gbps (Tables S1 and S2). Finally, Flye resulted in a more contiguous assembly with higher N50 and lower L50 in comparison to Canu for both 3 and 6 Gbps datasets (Fig. 3). Remarkably, Flye lead to the assembly of complete bacterial genomes in a range of only 2 to 21 contigs (Fig. S3).

### Assembly accuracy

Sequencing errors are the biggest throwback of third generation sequencing platforms. These errors can reach the final assemblies, resulting in lower quality draft genomes. In order to evaluate how the different assembles handle the MinION specific error profile, we ascertained the total number of SNPs and INDELs present in each draft metagenome. As described in the Methods section, we used two different -and complementary-strategies to quantify these type of errors: (1) minimap2 + bcftools, and (2) MuMMer (Fig. 4). Both strategies relied on the alignment of the draft assemblies to the reference metagenome, composed by a mix of all the complete genomes of each strain present in the mock community.

**Figure 4.**
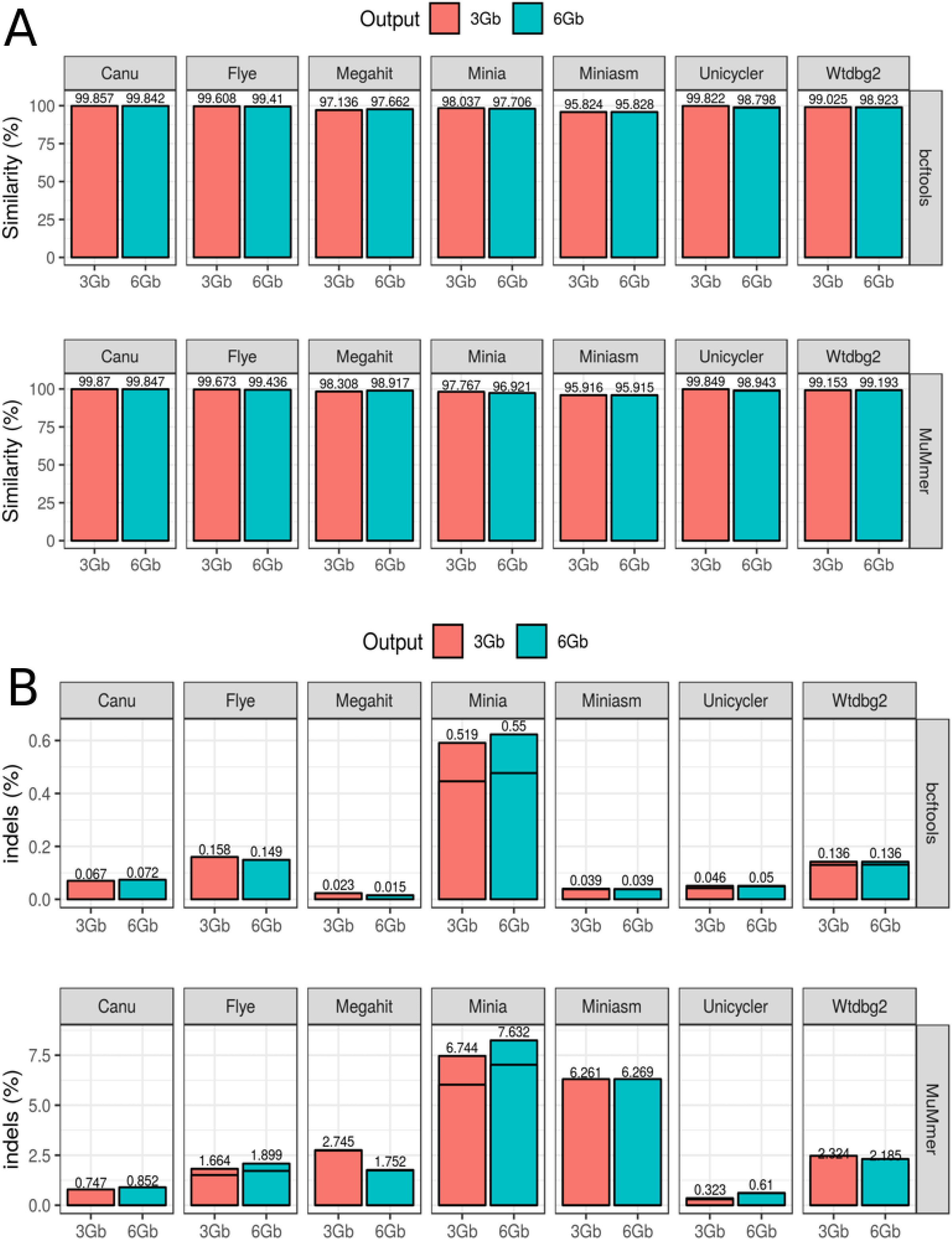
Assembly accuracy for the draft assemblies in the Even datasets. (A) Percentage of similarity calculated as the total number of matches normalized by the metagenome size; (B) Percentage of INDELs calculated as the total number of INDELs normalized by the metagenome size. In both cases, two different strategies were used: (1) alignment with minimap and evaluation with bcftools + ‘indels_and_snps.py’ in-house script; (2) alignment with MuMMer and evaluation with ‘count_SNPS_indels.pl’ script from Goldstein *et al*. (2019).

Results were not fully consistent between the two methodologies, especially for the INDELs estimation, but they still showed interesting trends. All the long-reads assemblers retrieved draft metagenomes with an average similarity higher than 98.9 %, with the exception of Miniasm, which resulted in an approximate accuracy of only 96%. Canu was the most accurate assembler for both methodologies and datasets, followed by Unicycler for the 3 Gbps dataset and Flye for the 6 Gbps one. In the case of the INDELs profile, Unicycler and Canu clearly outperformed Flye. Indeed, taking into account the lack of consistency of Miniasm results, Unicycler presented the lowest INDEL ratio. This might be explained by the polishing step via Racon (https://github.com/isovic/racon) that Unicycler pipeline incorporates. In order to test this hypothesis, we used Racon for polishing Flye assemblies with the original nanopore raw reads. In this case, no improvements were detected in SNPs and INDELs ratio.

### Biosynthetic gene cluster prediction

Gene prediction is highly affected by genome assembly and accuracy. Biosynthetic gene clusters (BCGs) are especially influenced by these factors, since they are usually found on repetitive regions which are often poorly assembled. In order to evaluate the BGC prediction on nanopore-based metagenomic assemblies, we used AntiSMASH to assess the number of clusters found on the draft assemblies retrieved by each tool in comparison to the reference metagenome (Fig. 5). For the 3 Gbps GridION dataset, Unicycler predicted the maximum number of BCGs (39/46), followed by Canu and Flye (38/46). Nevertheless, Flye BGC profile differed more from the reference profile, due to an enrichment in lasso peptides. To further study this phenomenon, lasso peptides predicted by Flye were searched though BLAST against the BGCs predicted in the reference metagenome. No hits were found, suggesting that these results might be assembly artifacts. For the 6 Gbps GridION dataset, Canu performed the best, but did not increase the number of predicted clusters (38/46). As expected, Unicycler drastically decreased the number of predicted BGCs. Interestingly, Flye performed worse with higher coverage, and resulted in less BGCs (32/46).

**Figure 5.**
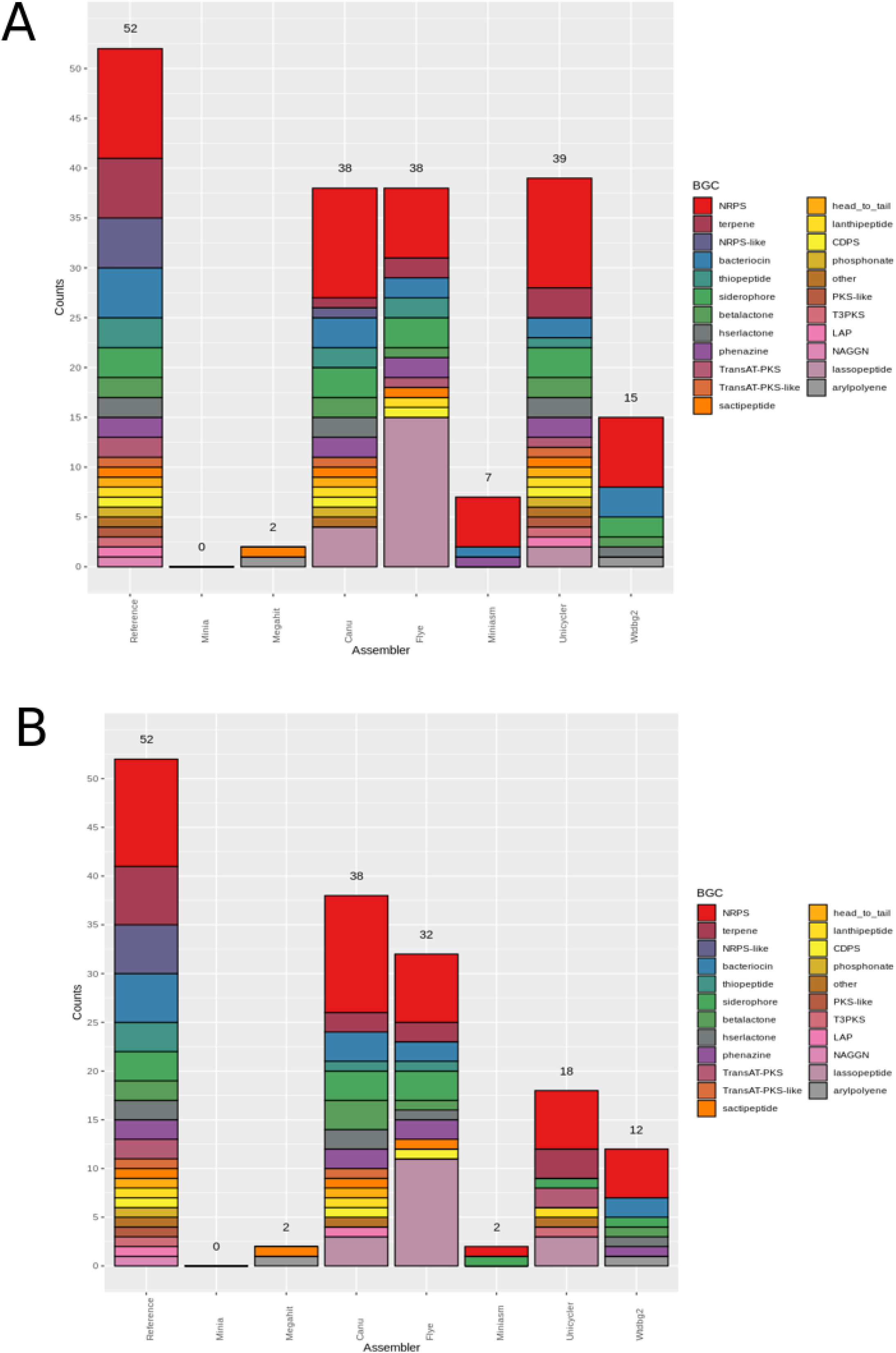
Number of biosynthetic gene clusters (BGCs) predicted by antiSMASH for each draft assembly in the Even GridION datasets. (A) BGCs predicted for the 3 Gbps dataset; (B) BGCs predicted for the 6 Gbps dataset.

## DISCUSSION

Assembling shotgun sequencing data is often a key factor for characterizing the functional and taxonomic diversity of microbial communities. In the recent years, MinION (Oxford Nanopore Technologies) sequencer is rapidly growing in popularity due to four basic reasons: (1) low cost, (2) long-reads generation, (3) portability, and (4) real-time analysis. Different bioinformatic tools have been developed in order to handle MinION sequences during the assembly process. Nevertheless, there is a lack of studies evaluating the performance of the current available tools for carrying out metagenomic assemblies from MinION sequences. This work aimed at filling this gap using data previously published by Nicholls *et al*. (2019), which consisted of the ultra-deep sequencing of two different mock communities (Table 1) using GridION and PromethION platforms (ONT). These sequencers follow the same sequencing principles than MinION, but they have a significantly higher output. For that reason, we decided to subsample the datasets to adequate their output to the current yield offered by MinION (3-6 Gbps) (Goldstein *et al*., 2019; Dhar *et al*., 2019; Parajuli *et al*., 2019).

Despite the relatively low complexity of the mock communities analyzed in this evaluation study, our results showed that there is a huge variation in assembly results depending on the software chosen to perform the analysis. Minia and Megahit poorly reconstructed the microbial genomes (Fig. 1 and Fig. 2) and produced highly fragmented draft assemblies (Fig. 3). This output was expected, since these assemblers are highly optimized to work on short-reads, which are totally different from the data generated by MinION.

Long-reads assemblers (Canu, Flye, Unicycler, Miniasm and wtdgb2) also presented significant differences in the general assembly performance. Overall, only Canu and Flye performed well on all the datasets tested. They were able to recover the eight bacterial genomes from the Even dataset with a high degree of completeness, and also reconstructed a significant fraction of the yeast genomes. Strikingly, the draft bacterial genomes were highly contiguous. In fact, Flye was able to reconstruct all the prokaryotic genomes in a range of only 2-21 contigs (Fig. S3).

Although sequencing errors are one of the main throwbacks of third generation data, Canu and Flye assemblies demonstrated to be up to 99.67% (Flye) and 99.87% (Canu) accurate. Regarding INDELs, Flye was more prone to insertion/deletions than Canu. This might influence the prediction of biosynthetic gene clusters, where Canu showed a more similar functional profile in comparison to the reference metagenome. Indeed, Flye BGC profile was biased to lasso peptides. BLAST analyses confirmed that these clusters did not match any other cluster predicted in the reference genome. This suggests that predicted lasso peptides might be artifact probably caused by frameshift erros due to INDELs, which explains that these type of cluster were more frequently detected in Flye’s assemblies -which had a higher INDEL ratio. Finally, time is a crucial parameter when choosing a bioinformatic tool, even more if considering MinION’s ability to generate real-time data. In this sense, Flye was up to 6.7 times faster than Canu, which resulted to be the slowest tool tested on this benchmarking.

Unicycler, miniasm and wtdbg2 results indicated that they are not suitable for metagenomic assembly due to different reasons. Unicycler worked well on the 3 Gbps Even dataset, but not for the rest. Indeed, this assembler was unable to run with the two 3 Gbps Log datasets, indicating a lack of consistency of the software for its application in a metagenomic context. Wtdbg2 was the fastest tool, but it was able to reconstruct only one complete genome for the Even datasets. For the Log datasets, wtdbg2 managed to recover the two most abundant bacterial genomes, being only outperformed by Canu and Flye. This fact suggested that the performance of wtdbg2 is associated with the composition of the original microbiome. Lastly, Miniasm resulted in low accuracy assemblies (~96 % of similarity to reference metagenome) (Fig. 4). This high error may explain the fact that metaQUAST failed to analyze Miniasm results. MetaQUAST is a tool mainly designed to work on second generation assemblies, and this error-prone assembly could have caused a problem when aligning the contigs against the reference. In fact, Miniasm’s low accuracy could be also detected in the prediction of biosynthetic gene clusters (Fig. 5). For the 3 Gbps dataset, antiSMASH was able to predict only 7 BGCs in the Miniasm assembly, whereas 15 BGCs were predicted in wtdbg2 assembly, despite having a lower metagenome recovery fraction (~42% in Miniasm vs. ~25% in wtdbg2).

To sum up, MinION data can lead to highly contiguous and accurate assemblies when using the proper tools, with no need of complementary sequencing with Illumina. From all the tested softwares, Flye resulted the best in terms of metagenome recovery fraction, metagenome size, and contiguity. Canu was the most accurate, introduced less INDELs, and resulted in a more similar BGC profile in comparison to the reference metagenome, but its assembly process also demonstrated to be time consuming. This work might help software developers to design new bioinformatic tools optimized for MinION-based shotgun metagenomic sequencing. Further research is still needed in order to evaluate the suitability of MinION for the metagenomic analysis of more complex microbial communities.

## CONCLUSIONS

Shotgun metagenomic sequencing based on short reads usually results in highly fragmented metagenomes, which complicate downstream analyses such as the recovery of individual genomes, or the prediction of complex and repetitive gene structures (i.e. biosynthetic gene clusters, CRISPR-CAS systems, etc). This work demonstrates that, despite the high error intrinsic to third-generation sequencing platforms, MinION sequencing alone can overcome these limitations and retrieve extremely contiguous genomes directly from simple microbial communities,. However, there is a huge variation in assembly performance depending on the chosen software. In general terms, Flye is the best assembler for MinION metagenomic data. This tool leads to the highest metagenome recovery ratio and performs robustly among the tested datasets. Canu is more suitable when lower error rates are required, as in the case of BGC prediction. Our results, along with the fast improvements of Oxford Nanopore devices and dedicated softwares, suggest that this type of platforms could become the metagenomic sequencing standard in the near future.

---

**Figure S1.** Fraction of genome covered by draft assemblies obtained with each tool, and for each individual microorganism (Log datasets). Minimap2 + miniasm assemblies are not shown, since it was not possible to evaluate them with metaQUAST.

**Figure S2.** General assembly performance of each tool for the Log datasets. (A) Run time; (B) N50; (C) Number of contigs; (D) L50.

**Figure S3.** Number of contigs for each bacterial genome retrieved by Flye for the Even datasets.

**Table S1.** Canu’s basic assembly statistics for the GridION datasets.

**Table S2.** Flye’s basic assembly statistics for the GridION datasets.

