## Supplementary material for "Assembly methods for nanopore-based metagenomic sequencing: a comparative study": Table S1

| | 3Gb | | | 6Gb | | |
|---|---|---|---|---|---|---|
| | **Contigs** | **N50** | **L50** | **Contigs** | **N50** | **L50** |
| **Bacillus subtilis** | 34 | 298,071.0 | 5 | 17 | 655,353.0 | 3 |
| **Enterococcus faecalis** | 13 | 388,478.0 | 3 | 10 | 747,976.0 | 3 |
| **Escherichia coli** | 5 | 2,669,962.0 | 1 | 6 | 4,941,166.0 | 1 |
| **Lactobacillus fermentum** | 13 | 402,806.0 | 2 | 14 | 4,941,166.0 | 1 |
| **Listeria monocytogenes** | 18 | 4,942,769.0 | 1 | 14 | 2,747,940.0 | 2 |
| **Pseudomonas aeruginosa** | 3 | 5,593,153.0 | 1 | 4 | 2,747,940.0 | 2 |
| **Salmonella enterica** | 3 | 4,942,769.0 | 1 | 11 | 2,075,612.0 | 2 |
| **Staphylococcus aureus** | 17 | 769,443.0 | 2 | 17 | 640,396.0 | 3 |
