## Supplementary material for "Assembly methods for nanopore-based metagenomic sequencing: a comparative study": Table S2

| | 3Gb | | | 6Gb | | |
|---|---|---|---|---|---|---|
| | **Contigs** | **N50** | **L50** | **Contigs** | **N50** | **L50** |
| **Bacillus subtilis** | 8 | 4,121,094.0 | 1 | 15 | 4,118,946.0 | 1 |
| **Enterococcus faecalis** | 4 | 2,916,346.0 | 2 | 8 | 2,917,992.0 | 1 |
| **Escherichia coli** | 12 | 686,395.0 | 2 | 19 | 1,282,129.0 | 2 |
| **Lactobacillus fermentum** | 6 | 1,916,903.0 | 1 | 21 | 1,909,954.0 | 1 |
| **Listeria monocytogenes** | 7 | 2,990,853.0 | 1 | 13 | 2,129,318.0 | 1 |
| **Pseudomonas aeruginosa** | 2 | 6,821,368.0 | 1 | 4 | 6,820,234.0 | 1 |
| **Salmonella enterica** | 12 | 1,848,158.0 | 2 | 15 | 1,282,129.0 | 2 |
| **Staphylococcus aureus** | 7 | 2,783,558.0 | 1 | 12 | 2,785,527.0 | 1 |
