## Supplementary figures and images for "Assembly methods for nanopore-based metagenomic sequencing: a comparative study"

### Figure S1

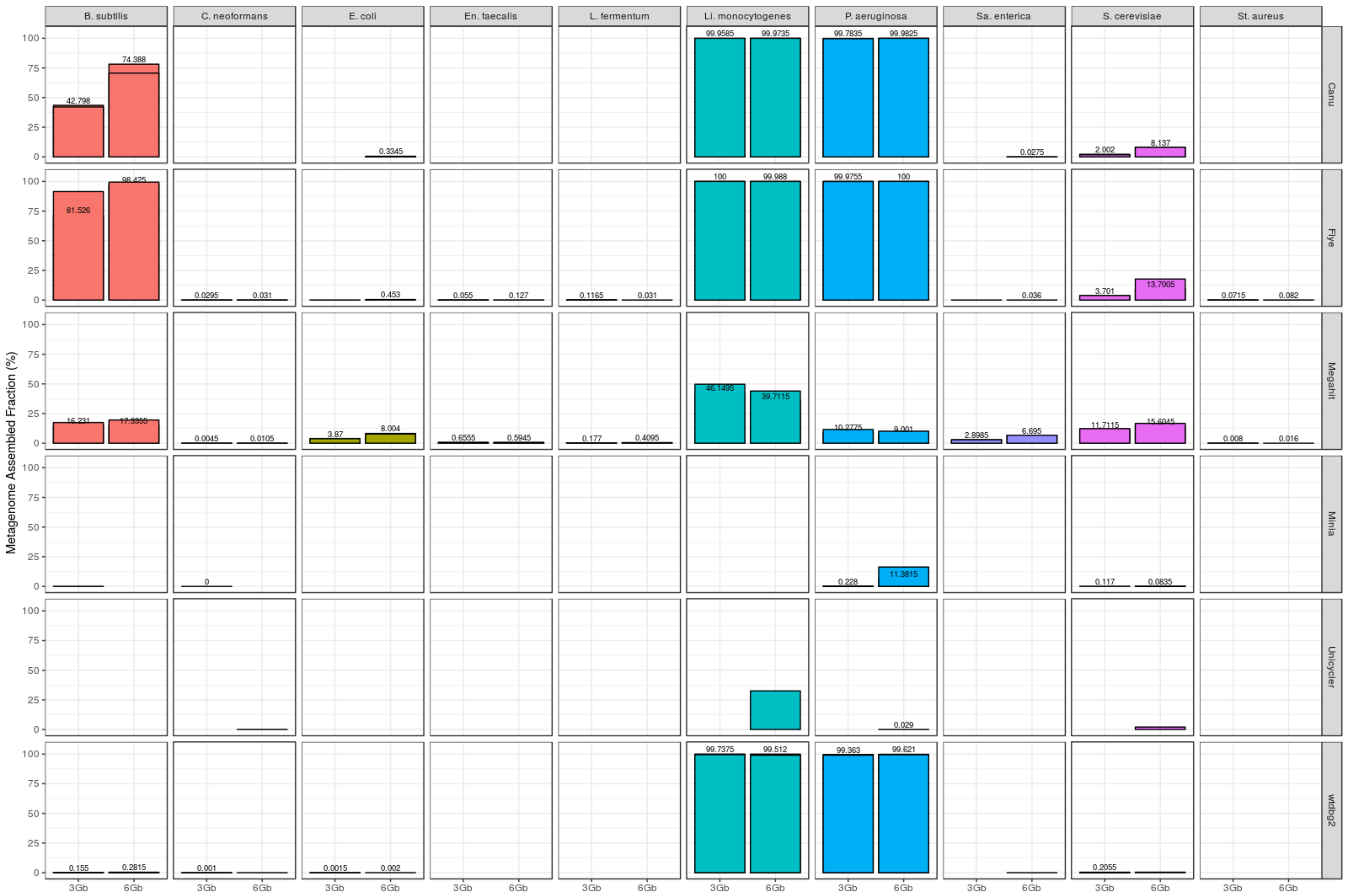

### Figure S2

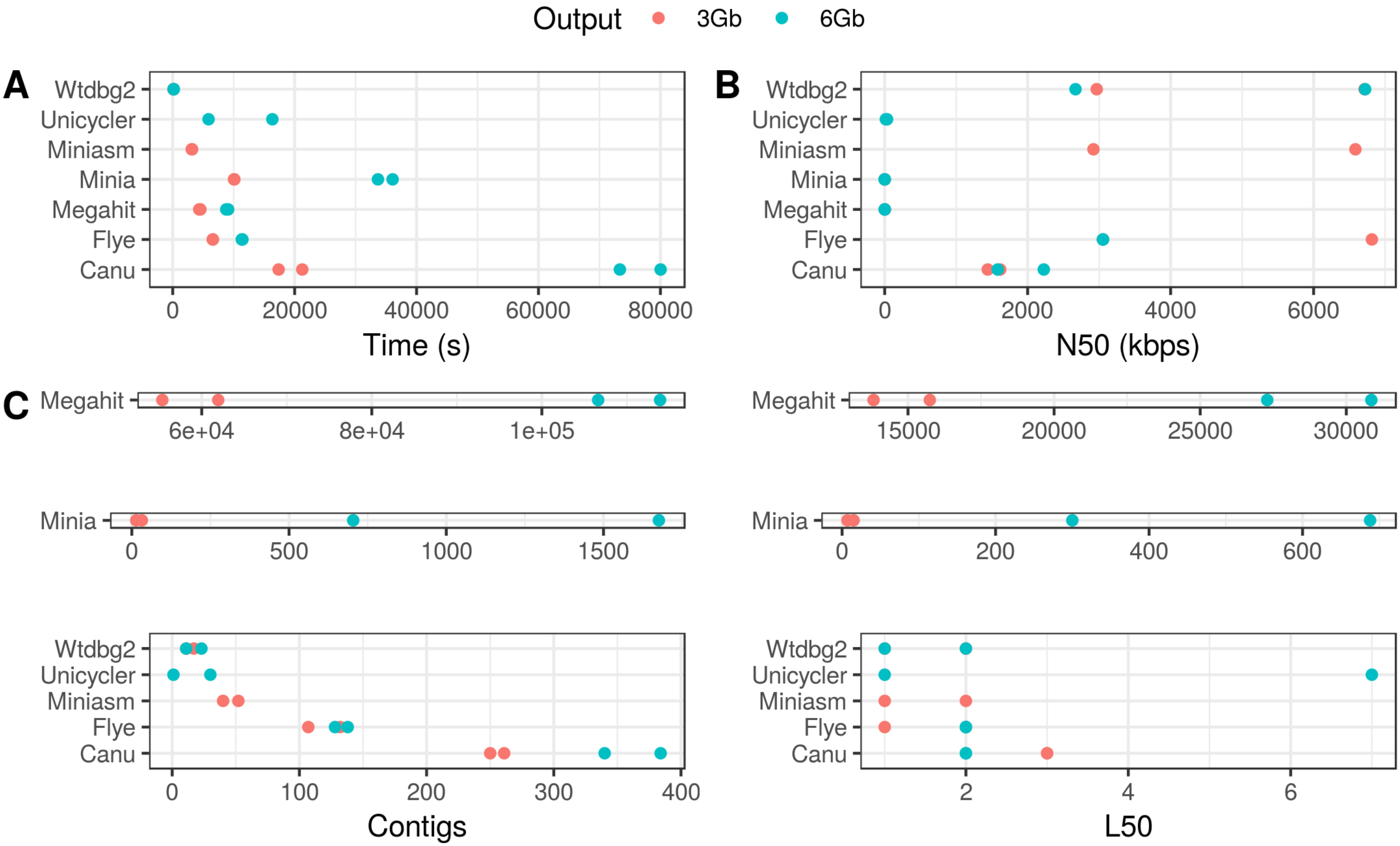

### Figure S3

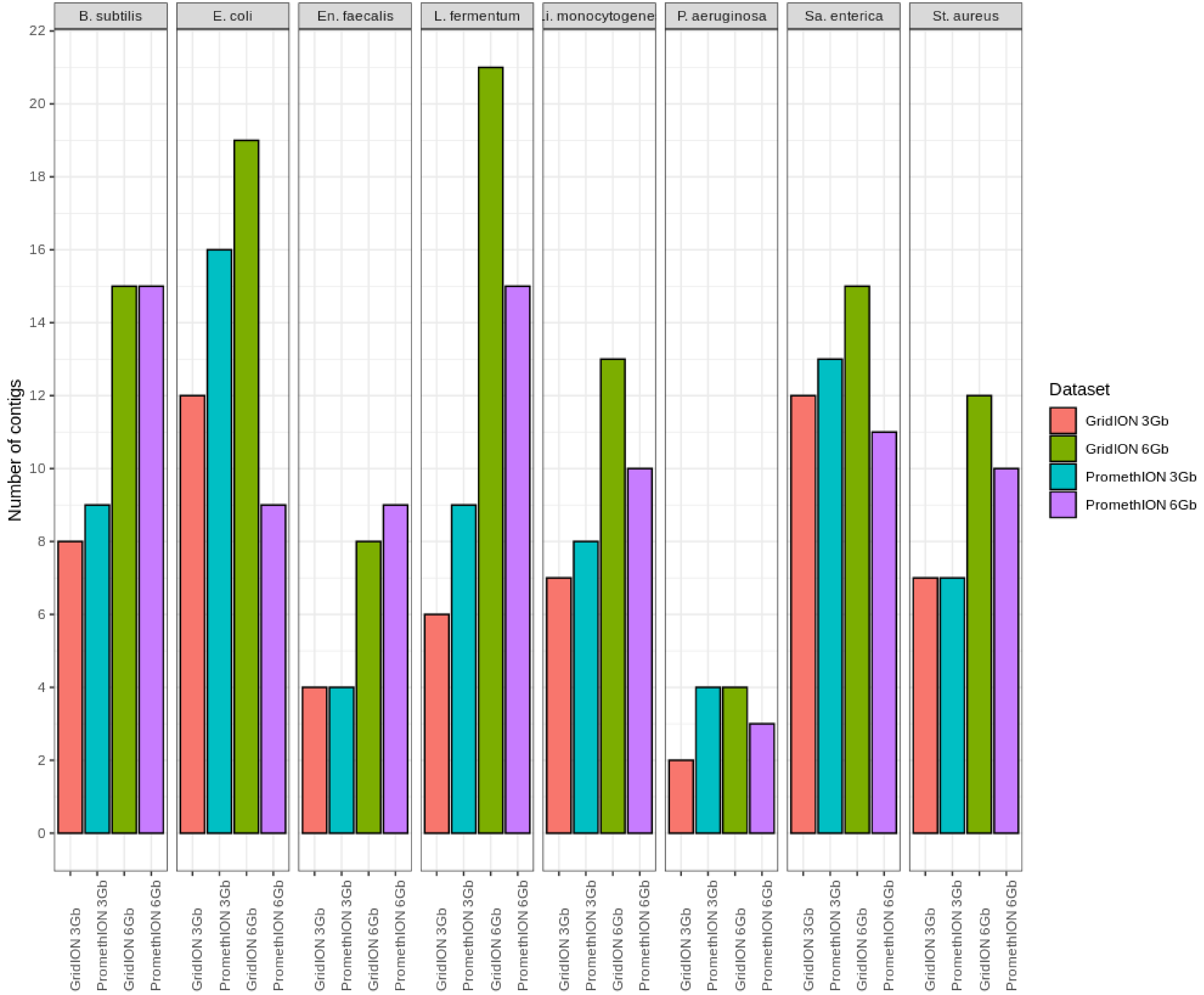
