## Supplementary File 1 for "Assembly methods for nanopore-based metagenomic sequencing: a comparative study"

#!/usr/bin/env python3
# -*- coding: utf-8 -*-
"""
Python script which print on the screen the number of SNPs and INDELs contained in a VCF file given as an argument.

Created on Mon Apr 15 12:37:15 2019

@author: Pascual Villalba-Bermell - Darwin Bioprospecting Excellence S.L.
"""
# Import module.
import sys

# Main program.
def main():
    # VCF file path.
    name_file = sys.argv[1]
    # Open VCF file.
    file_in = open(name_file, 'r')
    
    type_variants = []
    number_variants = []
    
    # Sum the SNPs and INDELs according to column number eight (INFO).
    for line in file_in:
        line = line.strip()
        
        if line[:2] != '##':
            data = line.split('\t')
            INFO = data[7].split(';')
            if INFO[0][:3] == 'DP=' and 'SNP' not in type_variants:
                type_variants.append('SNP')
                number_variants.append(1)
            elif INFO[0][:3] == 'DP=' and 'SNP' in type_variants:
                i = type_variants.index('SNP')
                number_variants[i] += 1
            elif INFO[0][:5] == 'INDEL' and 'INDEL' not in type_variants:
                type_variants.append('INDEL')
                number_variants.append(1)
            elif INFO[0][:5] == 'INDEL' and 'INDEL' in type_variants:
                i = type_variants.index('INDEL')
                number_variants[i] += 1
    
    # Print on the screen the number of SNPs and INDELs.
    print('\n\n## <type of variant>:<number of variants of this type>')
    for x in range(len(type_variants)):
        print(type_variants[x] + ':' + str(number_variants[x]))    


if __name__ == '__main__':
    main()
                
                
